# Development of an *in vitro* Experimental Model for Investigating the Effect of Matrix Stiffness on Epithelial Barrier Permeability

**DOI:** 10.1101/828079

**Authors:** Neilloy Roy, Emily Turner-Brannen, Adrian R West

## Abstract

Epithelial cells are well-known to be modulated by extracellular mechanical factors including substrate stiffness. However, the effect of substrate stiffness on an epithelial cell’s principal function –creating an effective barrier to protect the underlying tissue – cannot be directly measured using existing experimental techniques. We developed a strategy involving ethylenediamine aminolysis and glutaraldehyde crosslinking to chemically graft polyacrylamide hydrogels with tunable stiffness to PET Transwell membranes. Grafting success was evaluated using spectroscopic methods, scrape tests, and extended incubation in culture. By assessing apical to basolateral transfer of fluorescent tracers, we demonstrated that our model is permeable to biologically relevant molecules and usable for direct measurement of barrier function by calculating paracellular permeability.

We found that BEAS-2B epithelial cells form a more effective barrier on stiff substrates, likely attributable to increased cell spreading. We also observed barrier impairment after treatment with transforming growth factor beta, indicating loss of cell-cell junctions, validating our model’s ability to detect biologically relevant stimuli. Thus, we have created an experimental model that allows explicit measurement of epithelial barrier function for cells grown on different substrate stiffnesses. This model will be a valuable tool to study mechanical regulation of epithelial and endothelial barrier function in health and disease.

## INTRODUCTION

Cell structure and function are highly influenced by mechanical forces imparted by the extracellular environment. Transmission of forces onto the cytoskeleton by modalities such as stretch^1^, transmembrane compression^2,3^, shear flow^4^, and matrix stiffness^5^ initiate signaling cascades through the unfolding of protein domains and alterations in binding affinities at focal adhesions^6^ and cellular junctions^7^. Elucidation of these mechanotransduction pathways has been driven by the development of new technologies and experimental models that permit finely tuned cell mechanical challenges^8,9^. In particular, the creation of tunable polyacrylamide substrates bonded to glass coverslips^10^ have enabled experimental results demonstrating that extracellular matrix (ECM) stiffness directly affects cell phenotype^11–14^.

Epithelial and endothelial cells are mechanically regulated, with ECM stiffness known to modulate cytoskeletal dynamics^15,16^, proliferation^17^, transcriptional regulation^18^, epithelial-mesenchymal transition (EMT)^19–21^ and metastasis^22,23^. The principal function of these cell types is to separate luminal spaces from underlying tissues by forming cohesive barriers, with barrier function largely imparted by cell-cell adhesion mediated by cellular junctions^24–26^. Not only are these junction complexes the molecular basis of epithelial/endothelial barrier function, they are also hubs of mechanotransduction^8,27^. For example, collective migration during wound healing hinges upon cytoskeletal force transmission between adjacent cells through adherens junctions^28,29^, and at the same time, cellular junctions are downstream effectors of changes in cytoskeletal tension^30^. In the case of ECM stiffness, elasticity of the substrate at focal adhesions determines the amount of tension loaded onto the cytoskeleton and its force transducing elements^6^, suggesting an intimate connection between ECM stiffness and barrier function.

Explicit assessments of barrier function commonly include measurements of trans-epithelial/endothelial electrical resistance (TEER) and paracellular permeability^31^. Both these techniques require culturing cells on porous membranes (e.g., Transwells) made of polyester (polyethylene terephthalate; PET), polycarbonate or polytetrafluoroethylene (PTFE). Unlike materials such as polyacrylamide^10^, polydimethylsiloxane^32,33^, collagen^34^, alginate^23^ and other polymers that can be tuned to replicate ECM stiffness changes *in vitro*, the membrane stiffness of Transwell inserts presently cannot be adjusted.

To address this unmet need, we created a ‘Transwell-Substrate’ experimental model that allows ECM stiffness modulation of epithelial/endothelial cell monolayers and concomitant assessment of barrier function. Here we describe simple chemical modifications to commercially available PET Transwells that promote the adherence of polyacrylamide. These modifications were first demonstrated with spectroscopic and physical techniques on standard PET film and then integrated into a manufacturing protocol for Transwell-Substrates. We characterized and optimized substrate thickness and permeability before performing biological experiments in which epithelial cell permeability was assayed on different substrate stiffnesses under control conditions as well as after culturing with a known barrier antagonist, transforming growth factor beta (TGF-β). Our results demonstrate that epithelial permeability can be measured on Transwell-Substrates and that barrier function is impacted by underlying ECM stiffness, likely through a combination of altered cell spreading and modulation of cell-cell junctions. Transwell-Substrates are therefore a compelling experimental platform for assessing changes in epithelial/endothelial barrier function resulting from alterations in ECM stiffness.

## RESULTS

### Surface Chemistry of Transwell Membranes

Transwell membranes are a porous cellular growth surface that allow the formation of concentration and voltage gradients across a cell monolayer (Fig 1A). The chemical structure of PET is repeating units of terephthalate and ethylene glycol joined by ester linkages with the chemical formula (C_10_H_8_O_4_)_n_ (Fig 1B). The terephthalate portions of the overall structure consist of aromatic rings flanked by para-substituted carbonyl groups, which are part of the ester linkages. To confirm the chemical composition of PET Transwell membranes, the surface chemistry was analysed using Fourier Transform Infrared (FT-IR) spectroscopy against an ultrapure commercial PET film (Fig 1C). The prominent carbonyl (C=O; Fig 1C, i) stretching vibration was observed at 1714 cm^-1^ in Transwell membranes and at 1705 cm^-1^ in PET film. Aromatic vibrations (Fig 1C, ii) were seen at 1409 cm^-1^ and 1412 cm^-1^, 1022 cm^-1^ and 1022 cm^-1,^ and 871 cm^-1^ and 872 cm^-1^ in PET film and Transwells respectively, while ethylene bending vibrations (-CH_2_-; Fig 1C, iii) were observed at 1341 cm^-1^ and 1342 cm^-1^. Finally, the ester stretch (C-O; Fig 1C, iv) had four signals at 1232 cm^-1^, 1122 cm^-1^, 1092 cm^-1^ and 970 cm^-1^ for PET film and at 1247 cm^-1^, 1122 cm^-1^, 1089 cm^-1^ and 969 cm^-1^ for Transwell membranes.

**Figure 1.**
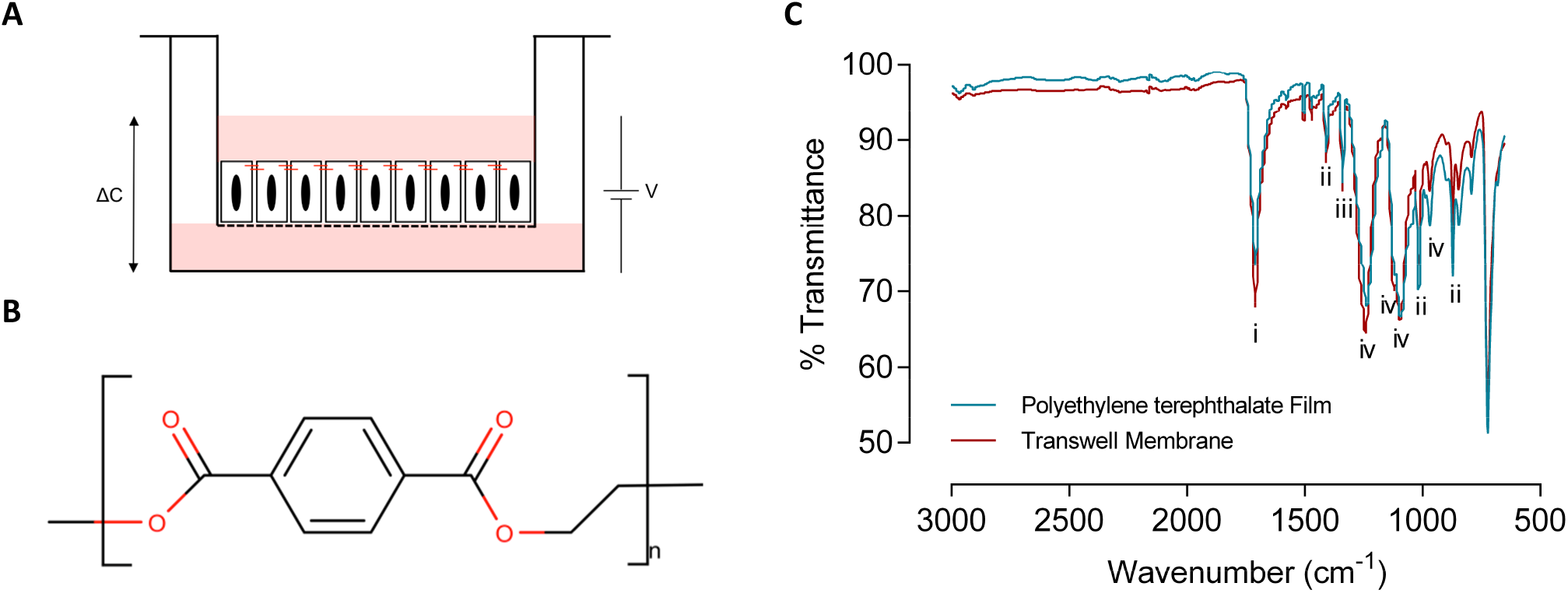
Transwell insert membranes are nominally composed of PET. (**A**) Schematic of a typical Transwell culture with a monolayer of epithelial cells. Cells are maintained on the porous PET membrane (dotted-line). The dual chamber design permits concentration gradients (ΔC) and potential differences (V) for explicit measurements of epithelial barrier function. (**B**) Chemical structure of PET. (**C**) FT-IR spectra of PET Transwell membrane and PET film. Spectra are annotated as follows: carbonyl (C=O) stretch (i), aromatic vibrations (ii) ethylene (-CH_2_-) bend (iii) and ester (C-O) stretch (iv). The spectra are highly similar indicating that Transwell membranes are indeed PET, albeit more crystalline than our ultrapure reference sample.

Collectively, these data indicate that Transwell membranes are indeed made of PET, with only minor discrepancies in spectra against our ultrapure reference sample. In particular, Transwells appear to have a stronger signal at 1122 cm^-1^ than PET film indicating a crystalline (C-O) stretch^35^. This is macroscopically evident when visually comparing Transwell membranes and PET film; crystalline-PET is opaque and amorphous-PET is transparent^36^. Nevertheless, the chemical likeness between PET Transwells and PET film allowed us to use the film to develop an effective strategy for grafting polyacrylamide to PET.

### Aminolysis of PET

We investigated and trialled numerous strategies to graft polyacrylamide to PET. Ultimately, we chose a strategy where PET film was chemically modified by ethylenediamine aminolysis (Fig. 2A) to create an amide (NH) bond (square) and surface amines (circle). The FT-IR spectrum of PET film after ethylenediamine treatment shows an amide (NH) bending vibration at 1540 cm^-1^ (Fig 2B, red) indicative of the amide linkage in the resulting PET-ethylenediamine conjugate. This amide signal is not present in the untreated PET film FT-IR spectrum (Fig 2B, blue).

**Figure 2.**
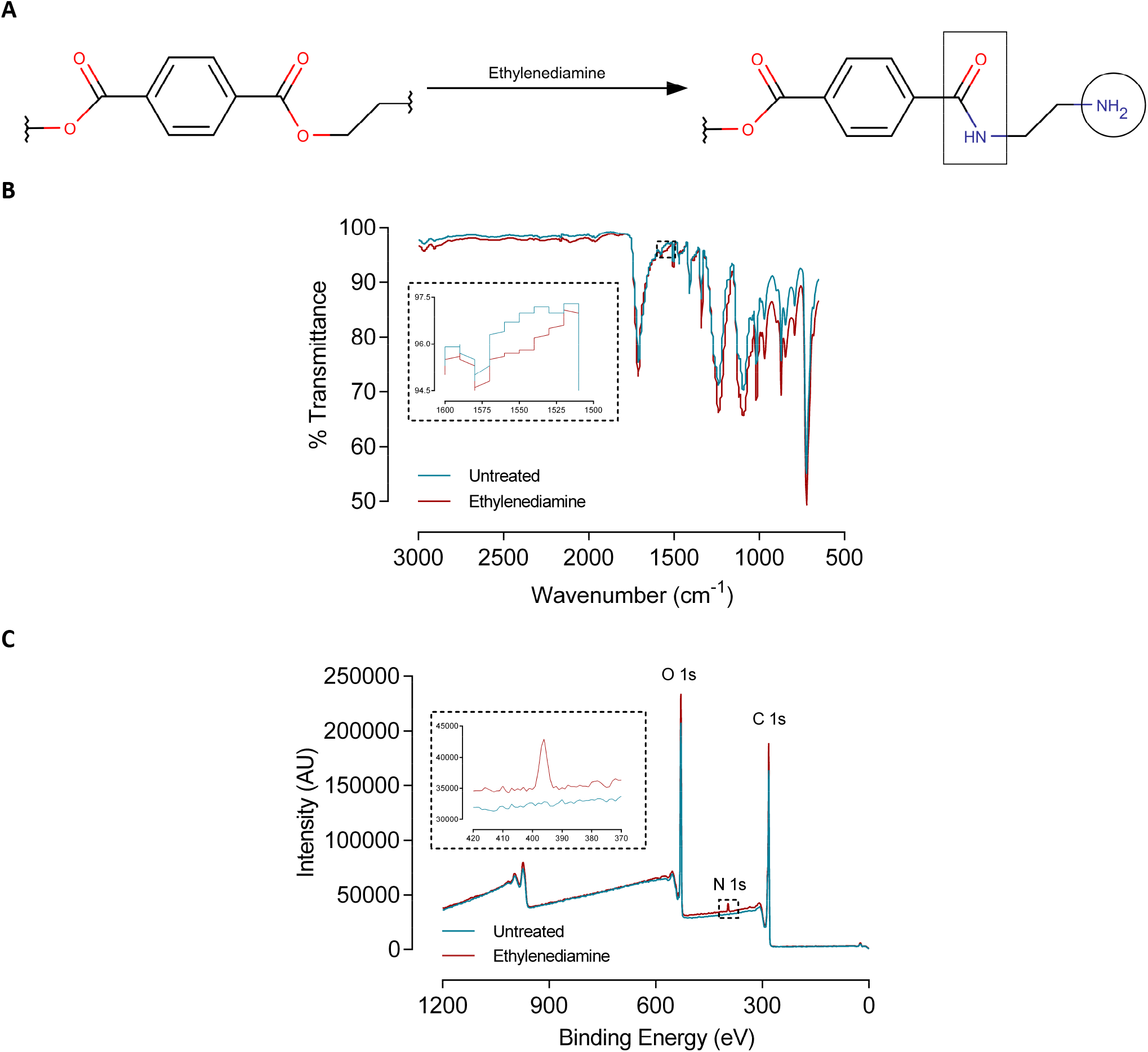
Ethylenediamine installs surface amines on PET through Aminolysis. (**A**) Aminolysis of PET by ethylenediamine. Amine groups attack PET carbonyls resulting in polymer chain scission. Dual amine functionality of ethylenediamine results in simultaneous amide linkage (square) and surface amine presentation (circle). (**B** and **C**) FT-IR and XPS survey spectra of untreated PET film and ethylenediamine treated PET film respectively. The peak at 1540 cm^-1^ (insert **B**) denotes an amide (NH) bending vibration and the peak at 396 eV (insert **C**) denotes nitrogen core electron binding energy, annotated as N 1s. Oxygen and carbon core electron binding energies are annotated as O 1s and C 1s respectively.

PET amination was confirmed with x-ray photoelectron spectroscopy (XPS). Both untreated PET film (Fig 2C, blue) and ethylenediamine treated PET film (Fig 2C, red) XPS survey spectra show strong signals at 529 eV and 281 eV, corresponding to oxygen (O 1s) and carbon (C 1s) core electrons respectively. Ethylenediamine treatment of PET film results in an additional peak at 396 eV (Fig. 2C, insert, red), which is indicative of nitrogen 1s electrons. Peak integration (Table 1) of the XPS survey data indicates the surface elemental composition (excluding hydrogen) of untreated PET film (75.5% carbon and 24.5% oxygen) agrees with the predicted composition of PET (71.4% carbon and 28.6%oxygen). With ethylenediamine treatment the surface composition of PET film changes to 74.1% carbon, 24.5% oxygen and, critically, 1.4% nitrogen.

**Table 1.** Surface elemental composition (%) of untreated and ethylenediamine-treated PET film. Predicted values were calculated according to the empirical formula of PET. The composition of treated and untreated PET film was obtained by integrating XPS survey spectra peaks.

| Element | Predicted PET | Untreated PET Film | Ethylenediamine treated PET Film |
| --- | --- | --- | --- |
| Carbon | 71.4 | 75.5 | 74.1 |
| Oxygen | 28.6 | 24.5 | 24.5 |
| Nitrogen | 0 | 0 | 1.4 |

### Grafting Polyacrylamide onto PET

Surface aminated PET film (Fig 3A, treated amine), was treated with glutaraldehyde, resulting in surface aldehyde groups (Fig 3A, treated aldehyde), which promoted fixation of polyacrylamide (Fig 3A, substrate). This was confirmed with qualitative scrape tests of different polyacrylamide substrates applied to ethylenediamine and glutaraldehyde treated PET. The scrape tests show polyacrylamide substrates applied to PET film treated with ethylenediamine and glutaraldehyde (Fig 3B, treated) were more difficult to remove than polyacrylamide applied to untreated PET film (Fig 3B, untreated), across the range of elastic moduli. Scrape tests, in conjunction with spectroscopic evidence, suggest ethylenediamine and glutaraldehyde can be used to graft polyacrylamide substrates with tunable stiffness to PET Transwells.

**Figure 3.**
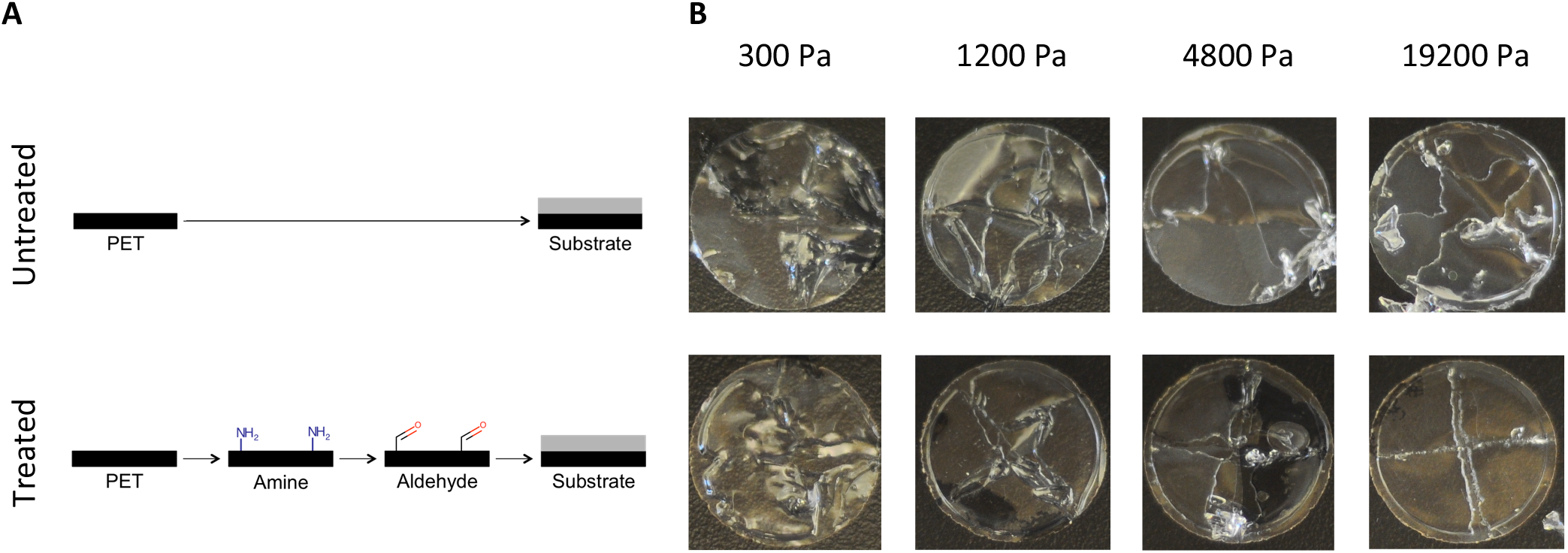
Ethylenediamine and glutaraldehyde treatment promote the adherence of polyacrylamide to PET. (**A**) Schematic of PET surface (black layer) modifications to achieve polyacrylamide (grey layer) adherence. Surface amines are installed by ethylenediamine treatment. Subsequent glutaraldehyde treatment results in surface aldehyde moieties. (**B**) Physical scrape tests of cured polyacrylamide resins applied to untreated and treated PET film. Our treatment regime makes it significantly more difficult to remove polyacrylamide from PET, particularly evident in the 19200 Pa sample. Treated PET maintains polyacrylamide adherence even after weeks in simulated cell culture conditions, whereas untreated film does not adhere well (not shown).

### Characterization of Polyacrylamide Applied to PET Transwell Membranes

The relationship between polyacrylamide resin volume, substrate stiffness, and substrate thickness was determined by casting different volumes of polyacrylamide in 24 mm diameter Transwells (Fig 4A, grey layer). Thickness of the resulting substrates was then measured (Fig 4B). As expected, substrate thickness is impacted by polyacrylamide resin volume (p<0.0001), but not by substrate stiffness (p=0.8574). These data demonstrate that polyacrylamide gels with different stiffnesses result in substrates with equivalent thicknesses at a given resin volume. Furthermore, a simple geometric relationship (i.e., volume of a cylinder) cannot be used to determine substrate thickness because the volume of polyacrylamide resin does not equal the volume of the polymerized substrate, presumably due to a small amount of resin loss through the base of the Transwell. Values were pooled across substrate stiffness to generate a calibration equation relating substrate thickness (*T*, μm) to polyacrylamide resin volume (*V*, μL) for a 24 mm Transwell (Equation. 1).

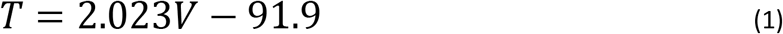

**Figure 4.**
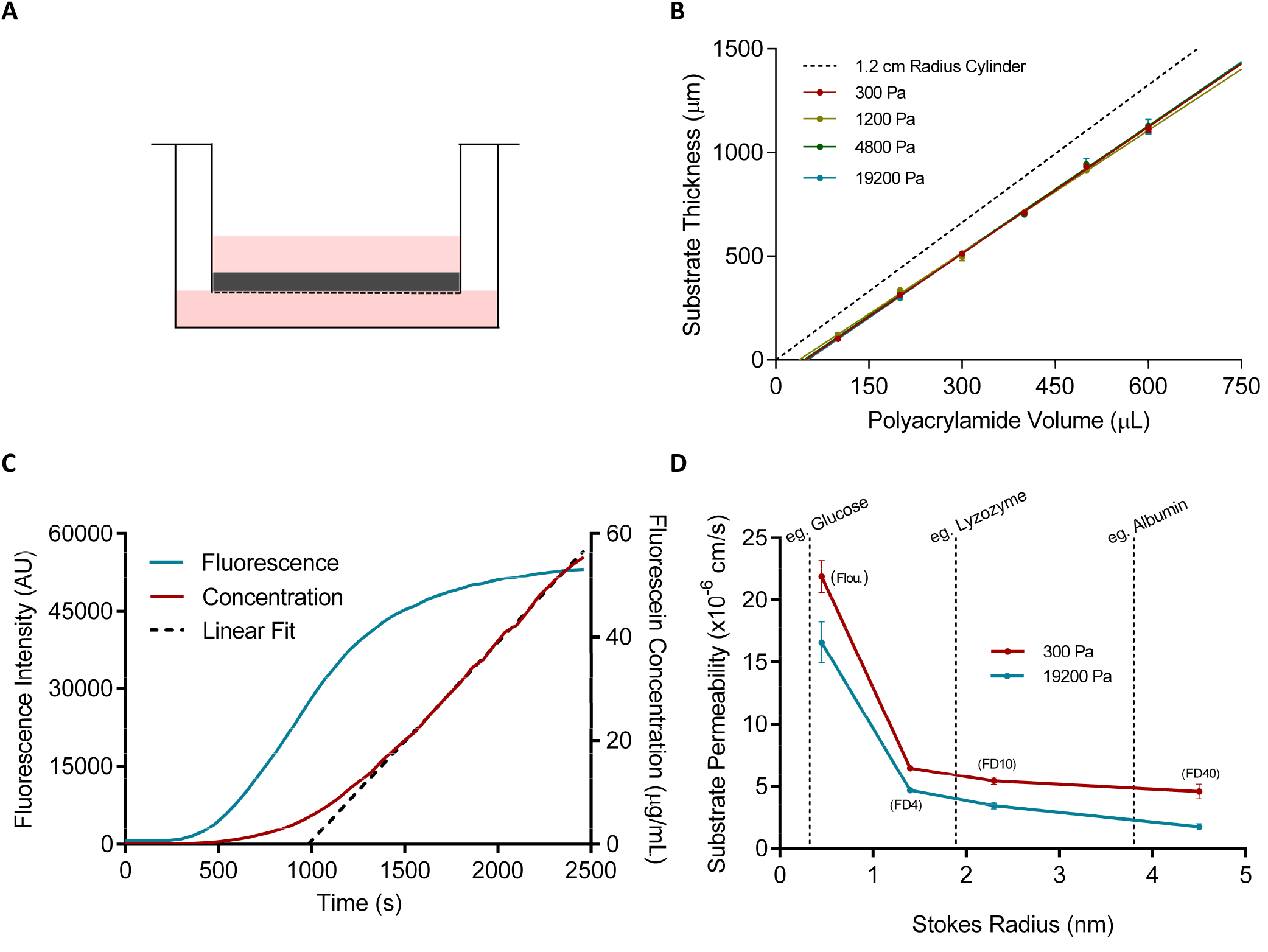
Polyacrylamide gels grafted to Transwell membranes permit the passage of molecules with varying sizes. (**A**) Transwell-Substrate diagram. Polyacrylamide (grey layer) is cast within a Transwell insert and grafted onto PET membranes (dotted line) modified with ethylenediamine and glutaraldehyde. (**B**) Thickness of polyacrylamide substrates cast in untreated Transwell inserts is linear with volume (R^2^>0.99 for all), and not dependent on stiffness (p<0.0001 for all, n=3). The dotted line shows the predicted relationship between resin volume and substrate thickness providing cured polyacrylamide perfectly assumes the geometry of the insert. (**C**) Basolateral chamber accumulation of fluorescein from the apical chamber diffusing through polyacrylamide (300 Pa, 919.5 μm thick) grafted to PET Transwell membrane. Fluorescence was monitored at 465 nm excitation and 520 nm emission wavelengths. The linear segment of the concentration-time curve is used to calculate permeability. (**D**) Permeability of 413.8 μm thick Transwell-Substrates measured with fluorescent tracers with different Stokes radii; both factors significantly affect permeability (Interaction p=0.1389, Stiffness p<0.0001, Radius p<0.0001, n=4). The dotted vertical lines represent the Stokes radii for commonly sized biological molecules, indicating compatibility of our model with standard cell culture.

Application of fluorescein to the apical chamber of Transwells and its subsequent accumulation in the basolateral chamber (Fig 4C, fluorescence) shows molecules are capable of diffusing through polyacrylamide grafted onto porous PET Transwell membranes. It is therefore possible to determine the absolute permeability of Transwell-Substrates by calculating the rate of apical-to-basolateral fluorescein accumulation (Fig 4C, concentration, linear fit). Substrate permeability is significantly impacted by substrate stiffness (p<0.0001) and Stokes radius (p<0.0001); soft 300 Pa substrates (Fig 4D, blue) are more permeable than stiff 19200 Pa substrates (Fig 4D, red) and tracer molecules with larger Stokes radii are less permeable than those with smaller Stokes radii. Stokes radius and substrate permeability do not interact (p=0.1389), indicating stiffness has a similar effect on permeability at all tracer sizes. Contrasting our tracers with compounds typically found in cell culture media suggests that our membrane permeability is biologically relevant.

### Epithelial Cell Permeability on Different Substrate Stiffnesses

To explicitly measure the isolated permeability of an epithelial cell layer (*P_E_*; cm/s) we must first measure the combined permeability of the substrate with cells (cellular; *P_CS_*, cm/s) and with cells removed (acellular; *P_AS_*, cm/s). PE can then be isolated from the substrate by calculating the reciprocal differences^37^ as per equation 2.

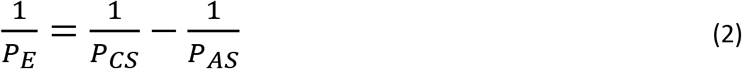

As expected, cellular Transwell-Substrates are less permeable than Transwell-Substrates from which the epithelial cells have been removed, irrespective of substrate stiffness (Fig 5A, each p<0.0001, n=6). Intriguingly, substrate stiffness had a significant effect on epithelial permeability (Fig 5B, p=0.0145, n=6), with cells on 300 Pa substrates (18.4 ±4.0 cm/s) forming a less effective barrier than cells on 2400 Pa (8.6 ±1.4 cm/s) and 19200 Pa (7.5 ±0.9 cm/s) substrates.

**Figure 5.**
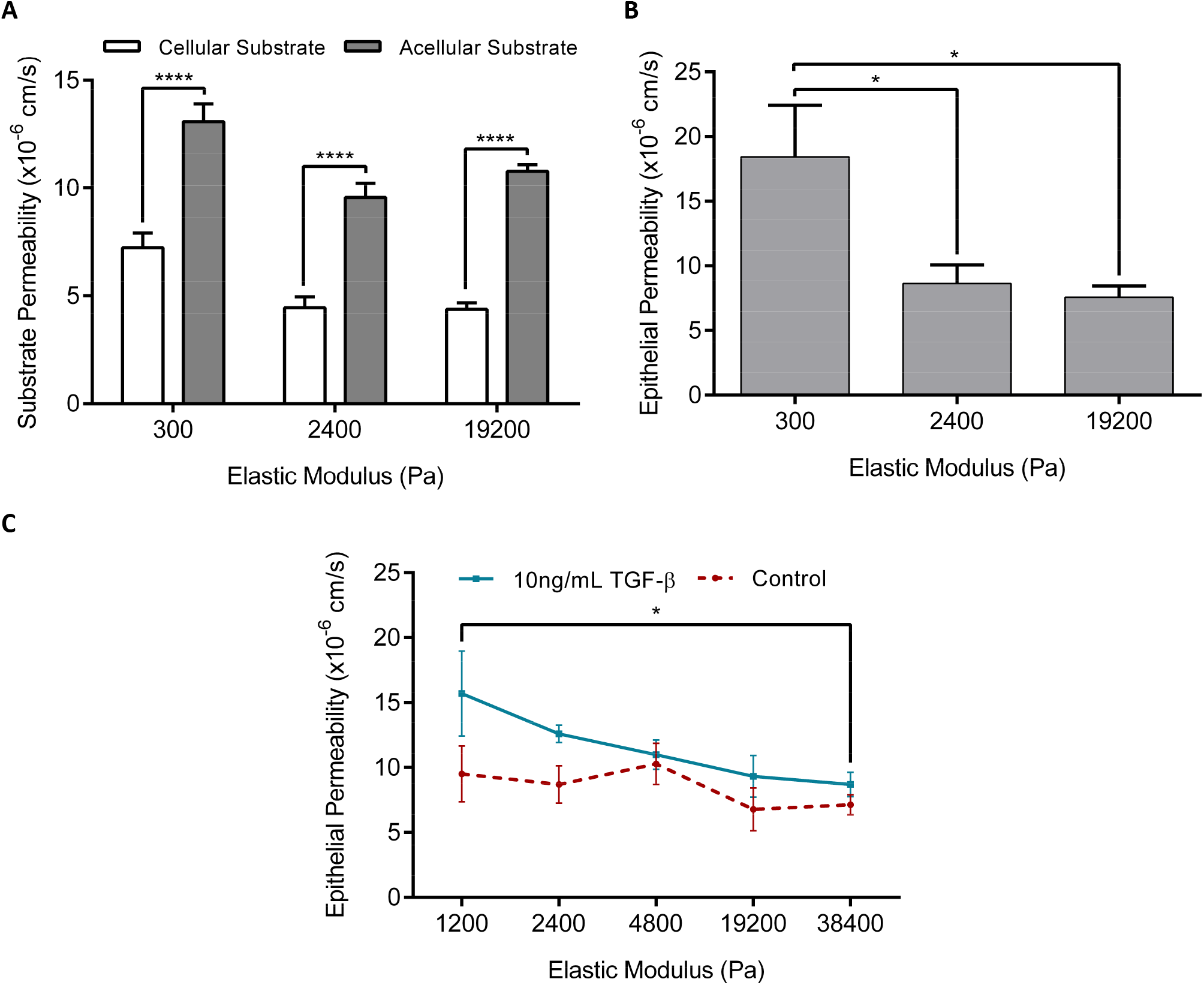
Substrate stiffness impacts permeability of BEAS-2B epithelial cells towards FD4. (**A**) Permeability of Transwell-Substrates measured 5 days after seeding with BEAS-2B epithelial cells (cellular), then remeasured after removing cells with trypsin (acellular). At each stiffness, cellular substrates were less permeable (p<0.0001 for all, n=6). (**B**) Calculated standalone permeability of the BEAS-2B epithelial layers grown on substrates with different elastic moduli. Cell on the stiffer substrates presented a more effective barrier (p=0.0145, n=6). (**C**) Cells across a range of substrate stiffnesses were dosed with TGF-β, and permeability was measured after 48 hours. Both stiffness and TGF-β significantly modulate barrier function (Interaction p=0.4947, Stiffness p=0.0347, TGF-β p=0.0068, n≥4).

To establish the ability of our model to measure biologically relevant changes in barrier function, epithelial cells were treated with transforming growth factor beta (TGF-β), which is a well-established antagonist of barrier function. As expected, TGF-β treated cultures were more permeable than untreated cultures across all stiffnesses, and the effect of stiffness was maintained in this experiment (Interaction p=0.4947, Stiffness p=0.0347, TGF-β p=0.0068, n≥4). Within the TGF-β treatment, cells on 1200 Pa (15.7 ±3.3 cm/s) substrates were more permeable than cells dosed with TGF-β on 38400 Pa (8.7 ±0.9 cm/s) substrates. Note that 300 Pa substrates needed to be excluded from this experiment, as TGF-β treatment on this substrate caused loss of cell adherence, effectively presenting no barrier. This resulted in virtually no difference between PCS and PAS, meaning equation 2 reported nonsensical permeability values that were negative or impossibly high. This highlights a limitation of our model’s sensitivity to measure epithelial barrier function as epithelial permeability becomes very high.

## DISCUSSION

Epithelial and endothelial barriers are essential to protect the underlying tissue from the environment and to regulate secretion of bioactive molecules to the luminal spaces. Barrier dysfunction is associated with numerous disease features, including structural remodelling of the blood vessels and airways. For this reason, we sought to create a model where the ECM stiffness experienced by barrier-forming cells could be manipulated while also allowing explicit measurement of barrier function. By grafting polyacrylamide substrates with a tunable stiffness to PET Transwells, we created the Transwell-Substrate model that enables simultaneous stiffness modulation and explicit measurement of paracellular permeability using the apical to basolateral diffusion of fluorescent tracers.

Chemically grafting a soft substrate to Transwells was necessary to ensure that it remains fixed during extended periods of time in cell culture. Polyacrylamide was chosen as the ideal soft substrate due to its common usage in mechanobiology research, established mechanical/chemical stability, and high porosity that would allow diffusion of waste and nutrient products. PET Transwells were chosen due to high transparency for imaging, and the range of options for chemical modification^38–45^. We also considered and trialed several other strategies to functionalise the PET including surface activation with oxygen plasma, silane treatment, and polydopamine coating, however these were discounted as either ineffective due to poor efficacy or undesirable due to the requirement for specialist equipment that may not be available in standard cell culture laboratories. Ultimately, we took inspiration from the original strategy of Pelham and Wang^10^, where 3-aminopropyltrimethoxysilane treated glass creates surface amines, and glutaraldehyde is used as a crosslinker to bond polyacrylamide. Although the chemistry of silica glass is distinct from PET, there are well-established methods for introducing amine groups onto PET, such as ammonia plasma^46^, ethylenediamine^47^ and polyvinylamine^48,49^. We chose ethylenediamine to install surface amines because it is liquid at room temperature, non-hazardous, and is easily rinsed away with methanol and/or ethanol.

Surface amine installation was confirmed by monitoring changes to the FT-IR spectra and XPS surveys of PET film after ethylenediamine treatment. The reaction of PET with ethylenediamine proceeds through a well-known mechanism called aminolysis^50^. Complete aminolysis depolymerizes PET into bis(2-aminoethyl) terephthalamide and α,ω-aminoligo (ethylene terephthalamide)^50^. However, partial aminolysis results in surface amine presentation through a PET-ethylenediamine surface adduct^47,51^ linked by an amide bond, which appears on FT-IR spectra as two signals^52^. Signal band I denotes the shifted amide carbonyl (N-C=O) stretches appearing on the shoulder of the ester carbonyl (O-C=O) stretch. This signal was not detected in our experiments as the more abundant ester carbonyl signal may have masked it. Signal band II denotes the amide (NH) bend, which was detected, suggesting PET aminolysis and possible amine functionality. XPS was performed in conjunction with FT-IR spectroscopy to support PET surface modification, with the emergence of a signal peak at 396 eV suggesting nitrogen incorporation following aminolysis. It should be noted that neither FT-IR nor XPS directly demonstrate amine functionality post ethylenediamine treatment which would need to be confirmed by other assays^48,53^ or spectroscopic techniques^54,55^.

Nevertheless, presumption of installed surface amines allowed us to proceed with glutaraldehyde treatment to install surface aldehyde moieties, which promote the adherence of polyacrylamide to the underlying scaffold as previously documented^5,56–58^. Spectroscopic analysis of PET film consecutively treated with ethylenediamine then glutaraldehyde was not performed as it is unlikely any spectral changes would be detectable due to the similarity of the chemical linkages between glutaraldehyde and PET. We can also say with confidence that glutaraldehyde will react with surface aminated PET because the Schiff Base reaction between amines and aldehydes are known to occur readily^59^. In lieu of quantitative analyses for surface aldehyde groups we performed qualitative scrape testes on polyacrylamide applied to ethylenediamine and glutaraldehyde treated PET film. We were able to demonstrate strong adhesion, indicating our surface modification strategy was effective.

In translating our grafting strategy from PET film to Transwells, we first discovered that the adhesive seal bonding the membrane to the insert can be dissolved by prolonged ethylenediamine exposure. We found that three repeated 90 second treatments with ethylenediamine interspersed with methanol washing preserves Transwell integrity and installed sufficient surface amines such that polyacrylamide adhered to treated PET for many weeks in simulated culture conditions.

Polyacrylamide is porous and its diffusion characteristics have previously been analyzed using fluorescent tracers^60^. We showed that a barrier composed of polyacrylamide and porous PET is indeed permeable by measuring the flux and accumulation of fluorescein from the apical chamber to the basolateral chamber. We characterized Transwell-Substrate permeability with a series of florescent tracers with Stokes radii between 0.45 nm and 4.5 nm to replicate the range of soluble biomolecules present in culture media such as glucose (0.32 nm)^61^ and albumin (3.8 nm)^62^. Molecules of different sizes can move between the two chambers indicating Transwell-Substrates should be feasible for cell culture and epithelial barrier function experiments.

Optimization of polyacrylamide resin volume and thickness was a critical step to ensure we could achieve both substrate permeability and biologically relevant mechanotransduction. Barrier thickness is inversely proportional to permeability^37^, therefore a thinner substrate is favorable for cell culturing because nutrients and waste can easily diffuse between both Transwell components. That said, very thin substrates are not desirable because as substrate thickness approaches the lateral dimensions of the cell the substrate is sensed as stiff regardless of actual substrate stiffness; threshold thicknesses have been reported as 60 μm for rat cardiofibroblasts^63^ and 100 μm for human fetal lung fibroblasts^64^. Ultimately, 140 μL of polyacrylamide resin was the minimum usable volume to reliably cover Transwell membranes and create a flat intact cellular layer. This corresponds to a substrate thickness of 191 μm, which is sufficient for appropriate mechanotransduction for the vast majority of cultured cells.

We initially attempted to use Calu-3 airway epithelial cells to demonstrate the utility of Transwell-Substrates in barrier function experiments due to their excellent barrier forming properties (baseline TEER ≥ 1600 Ω·cm^2^ and baseline FD4 permeability ≤0.006×10^-6^ cm/s^65–67^). However, we found that Calu-3 adhere poorly to softer substrates necessitating a switch to BEAS-2B cells which adequately adhered to substrates across the desired stiffness range. Although BEAS-2B cells are not typically used in epithelial barrier function studies, there are reports in which the TEER^68–71^ and permeability^72–74^ of BEAS-2B cells have been measured. Our results collectively show that ECM stiffening decreases BEAS-2B permeability and thus improves barrier function. This is perhaps in contrast to previous reports; Krishnan *et al*^30^ indirectly demonstrated endothelial barrier dysfunction on stiff ECM by observing similar cytoskeletal architecture to that observed during increased epithelial permeability. Huynh *et al*^75^ directly showed stiff ECM increases the relative permeability of endothelial cells as assessed by quantifying fluorescent tracer intensities in z-slice images of the substrate polyacrylamide beneath the endothelial layer.

A possible explanation for this discrepancy is that epithelial and endothelial cells may have a biphasic barrier function response to ECM stiffness. Mammoto *et al*^76^ demonstrated this biphasic response in blood vessels of *ex vivo* tissue where TEER peaks at a particular tissue stiffness. In this paradigm, cell spreading dominates at lower stiffnesses and promotes barrier function simply by virtue of cells occupying more area. At higher stiffnesses disruption of cell junctions by enhanced cytoskeletal contractility antagonizes barrier function. We are likely demonstrating the cell-spreading phase in our experiments by virtue of using the weakly-barrier-forming BEAS-2B cells; there may have been minimal intact epithelial junctions to disrupt, or insufficient cytoskeletal tension to induce any significant change.

Nevertheless, TGF-β increased permeability across the stiffness range. This is consistent with its known ability to downregulate epithelial characteristics including junction protein expression in BEAS-2B cells^77,78^. This would suggest at least some contribution from cell-cell junctions to barrier function, and their loss with TGF-β treatment. Alternatively, the increased permeability we measured may be attributable to TGF-β induced apoptosis, which has been reported to be enhanced on soft substrates^19^, and may explain our inability to record accurate permeability measurements from TGF-β treated cells on 300 Pa substrates. Testing alternative epithelial and endothelial cell types with our model, particularly those with higher barrier-forming capacity, will be required to fully understand the dynamics of barrier function across different substrate stiffnesses.

In summary, we have reported a workflow for chemical modification of commercially available PET Transwells using readily available equipment and reagents, to create an experimental model in which substrate stiffness can be modulated and absolute epithelial permeability measured. These Transwell-Substrates will be highly beneficial to explore the connection between matrix stiffness and barrier function, and for *in vitro* studies of diseases in which changes in ECM stiffness and barrier function are concomitant pathophysiological features.

## ACKNOWLEDGEMENTS

Adrian West is supported by NSERC Discovery Grant # RGPIN-2014-06412 and Research Manitoba New Investigator Operating Grant # 1653. Neilloy Roy is supported by a Research Manitoba and Children’s Hospital Research Institute of Manitoba Graduate Studentship (# 1329). The authors would like to thank Dr. Kevin McEleney from the Manitoba Institute for Materials for his assistance with spectroscopy.

## AUTHOR CONTRIBUTIONS

Neilloy Roy designed and performed experiments, analysed data and co-wrote the manuscript.

Emily Turner-Brannen performed cell culture and co-wrote the manuscript.

Adrian West conceived the study, designed experiments and co-wrote the manuscript.

## METHODS

### Polyethylene Terephthalate Film Surface Modification

PET film sheets (13 μm thick, Huntingdon 893-576-19) were punched into test slips with a 19 mm diameter hole-puncher (EK Tools 54-10054). PET slips were then washed with ethanol three times and air dried before being submerged in ethylenediamine (Sigma-Aldrich E26266) for 45 minutes in glass petri dishes. PET slips were washed with methanol and then ethanol three times and thoroughly dried before being treated with 1%glutaraldehyde (Sigma-Aldrich G7776) in phosphate buffered saline (PBS; Gibco 70011-044) for 1 hour. They were then washed three times with ddH_2_O and air dried.

### Spectroscopy

FT-IR spectra of Transwell membranes and PET film slips were collected with a ThermoNicolet Nexus 870 Infrared Spectrometer. A Kratos Axis Ultra X-ray Photoelectron Spectrometer was used to collect X-ray photoelectron spectra (XPS). XPS surveys peaks were integrated using CasaXPS software (Casa Software Ltd.).

### Polyacrylamide Scrape Test

One hundred microliters of polyacrylamide resin containing different ratios of 40% acrylamide (Bio-Rad 161-0140) and 2%bis-acrylamide (Bio-Rad 161-0142) were mixed with ammonium persulfate (APS; Bio-Rad 161-0700) and tetramethylethylenediamine (TEMED; Bio-Rad 161-0801) as described previously [Ref (^79^) and Table 2]. The solutions were immediately applied to control and ethylenediamine/glutaraldehyde treated PET film slips. Resins were then ‘sandwiched’ with 18 mm round glass coverslips (VWR 48382-042) coated with Rain-X (Permatex BCRX11212CN) and left to polymerize for 30 minutes. Coverslips were separated from the polyacrylamide and grafting success was assayed by scraping substrates with forceps. Adherence was qualitatively assessed, and results were photographed.

**Table 2.** Polyacrylamide recipes corresponding to different substrate stiffnesses. Polymerization was initiated by adding 500 μL of 1% APS in ddH_2_O and 7.5 μL TEMED.

| Elastic Modulus (Pa) | 40% Acrylamide ($\mu\text{L}$) | 2% Bis-acrylamide ($\mu\text{L}$) | Water ($\mu\text{L}$) |
| --- | --- | --- | --- |
| 300 | 375 | 121 | 3996.5 |
| 1200 | 375 | 268.5 | 3849 |
| 2400 | 937.5 | 84 | 3471 |
| 4800 | 937.5 | 133.5 | 3421.5 |
| 19200 | 937.5 | 591 | 2964 |
| 38400 | 1500 | 294 | 2698.5 |

### Substrate Thickness

Transwell inserts with 24 mm diameter and 0.4 μm pore size (Corning 3450) were placed on 75×38×1 mm glass microscope slides (Fisher 12-550B) coated with Rain-X. Various volumes of polyacrylamide resin (Table 2) were applied to Transwell membranes, and gels were cast by covering the resin with 22 mm round coverslips (Fisher 125461) coated with Rain-X. Gels were left to polymerize for 30 minutes before the inserts were separated from the microscope slide. With the coverslips left in place the combined thickness of the substrate and coverslip was measured using a digital micrometer (Fowler 548150012). The coverslip was then removed, and substrate thickness was calculated by subtracting the coverslip thickness from the total thickness.

### Cell Culture

BEAS-2B adenovirus 12-SV (simian vacuolating) 40 virus hybrid transformed human bronchial epithelial cells were purchased from ATCC (CRL-9609). Cells were grown in 75 cm^2^ tissue culture flasks coated with an epithelial coating solution (ECS) comprised of 0.01 mg/mL human fibronectin (Corning 354008), 0.03 mg/mL rat-tail collagen-I (Corning 354236) and 0.01 mg/mL bovine serum albumin (BSA; Sigma A9418) dissolved in LHC basal media (Gibco 12677027). Cells were seeded at 3×10^3^ cell/cm^2^ and maintained in LHC-9 media (ThermoFisher 12680013) at 37 °C with 85% humidity and 5% CO_2_. Once the cells reached 80-85% confluence they were briefly washed with PBS and treated for 10-20 minutes with TrypLE cell disassociation reagent (Gibco 12604039) at 37°C. Cells were then collected in 2 mL of soybean trypsin inhibitor (Sigma T6414), centrifuged (150 RCF, 5 min, 18°C) and re-suspended in LHC-9 media. Cells were counted and re-seeded for culturing or experimentation, with media changed after 24 hours then every 2 days.

### Transwell-Substrate Manufacturing

Transwells inserts were removed from their 6-well plates and placed into Pyrex petri dishes inside a fume hood. The PET membranes were then covered with ethylenediamine for 90 seconds and washed with methanol. This treatment/washing cycle was repeated a total of three times to generate sufficient surface amine installation without dissolving the adhesive bonding the Transwell membrane to the insert. After an ethanol wash and air drying, the inserts were treated with 1 mL of 1%glutaraldehyde in PBS for 45 minutes, washed three times with ddH_2_O and left to dry. Inserts were then placed onto glass microscope slides (75×38×1 mm) coated with Rain-X, 140 μL of polyacrylamide resin applied (Table 2), and substrates cast by immediately sandwiching with a Rain-X coated 22 mm round coverslip. After 30 minutes of curing, coverslips and slides were separated from the cast substrates.

Substrates were placed into 6-well plates and both chambers were washed three times with 50 mM HEPES pH 8.5 (Sigma H3375). The polyacrylamide was activated for ECM protein coating by covering the substrates with 500 μL of 0.5 mg/mL sulfo-SANPAH (Proteochem c1111) in HEPES buffer and exposing to 365 nm UV light for 6 minutes (Spectrolinker XL-1000, full power). Photoactivation was repeated once with a fresh aliquot of sulfo-SANPAH before substrates were washed three times with HEPES to remove unreacted reagent. ECS (1 mL) was added to aseptically to substrates and left for 24 hours at 4°C. Substrates were washed three times with sterile PBS, placed into new 6-well plates (Costar 720083) and UV sterilized for 20 minutes. BEAS-2B were seeded onto substrates at 200×10^3^ cells/cm^2^ and both chambers were made up to 2 mL of LHC-9 media.

### TGF-β treatment

Lyophilised TGF-β (StemRD TGFB1-005) was reconstituted at 1 mg/mL in dilution buffer composed of 0.1% BSA in 5 mM HCl. Four days post-seeding, media in both Transwell chambers was replaced with LHC-9 media containing an appropriate concentration of diluted TGF-β or equivalent volume of dilution buffer. Cells were maintained in TGF-β for 48 hours prior to permeability measurements.

### Transwell-Substrate Permeability

The permeability (*P*; cm/s) of Transwell-Substrates (surface area *A* = 4.67 cm^2^) was determined by measuring the diffusion of fluorescent tracers (Table 3) from the apical to basolateral chamber. For acellular experiments, the basolateral chamber was loaded with PBS (*V_B_* = 2 mL) and the apical chamber loaded with 2 mL of tracer (concentration *C_A_* up to 2000 μg/mL) dissolved in PBS. Immediately following apical chamber loading, a FLUOstar Optima fluorescence plate reader was used to continuously monitor tracer accumulation in the basolateral chamber every minute for 2 hours. Fluorescence-time curves for each substrate were converted to concentration-time curves using a fluorescent standard curve and the hyperbolic interpretation function of GraphPad Prism 6.07. The slope of the linear segment of these curves (*ΔC_B_*/*Δt*; μg/mL/s) was determined using the linear regression function of GraphPad. These slopes relate to permeability by the steady-state approximation for Transwells^31^ (Equation 3):

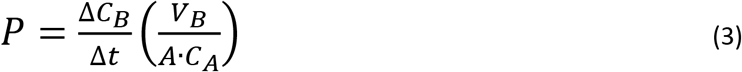

**Table 3.**
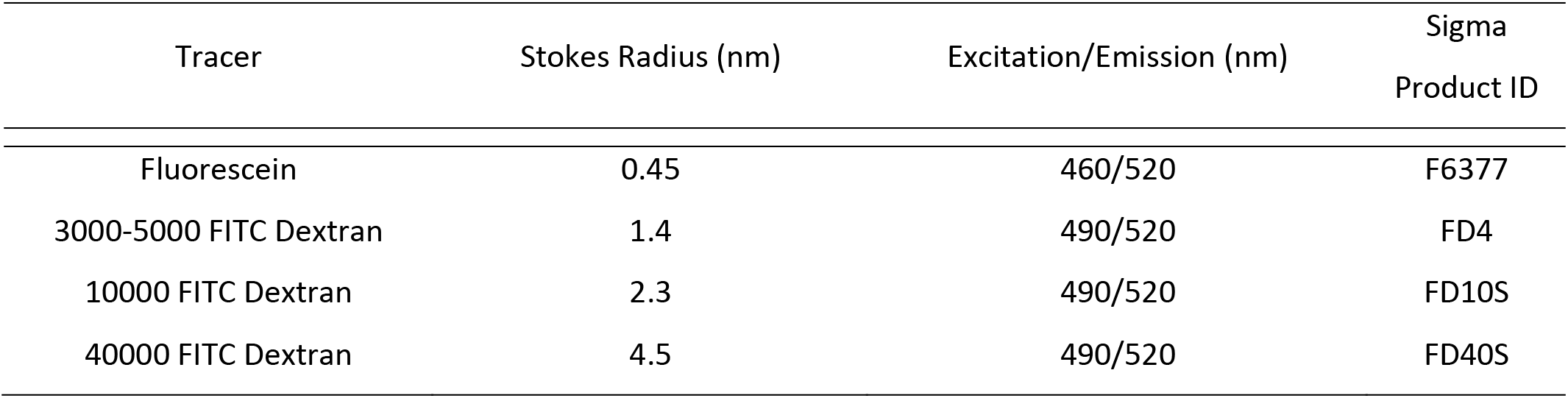
Physical parameters and measurement values for fluorescent tracers used in permeability experiments.

| Tracer | Stokes Radius (nm) | Excitation/Emission (nm) | Sigma<br>Product ID |
| --- | --- | --- | --- |
| Fluorescein | 0.45 | 460/520 | F6377 |
| 3000-5000 FITC Dextran | 1.4 | 490/520 | FD4 |
| 10000 FITC Dextran | 2.3 | 490/520 | FD10S |
| 40000 FITC Dextran | 4.5 | 490/520 | FD40S |

For experiments with cellular substrates media was removed from both chambers and they were washed once with sterile PBS. The apical chamber was loaded with 2 mL of filter-sterilised 1000 μg/mL FD4 dissolved in Hank’s Balanced Salt Solution (HBSS, Gibco 14065056). Assessment was initiated by transferring substrates to fresh 6-well plates containing 2 mL of sterile HBSS per well. Fluorescence readings were taken at 37°C and temperature was maintained at 37°C throughout the experiment.

### Transwell-Substrate Decellularization

After cellular substrate permeability values were measured, all tracers and HBSS were removed. Substrates were then loaded with 2 mL of 0.25% trypsin (HyClone SH3004201) and placed in an incubator for at least 30 minutes. Substrates were thoroughly washed at least three times with PBS to remove any and all traces of cellular debris, then remeasured for acellular permeability.

### Data Analysis and Statistics

All numerical data are presented as mean ± standard error. Statistical tests were performed with the GraphPad Prism 6.07 software package, with p<0.05 considered statistically significant. 2-way ANOVA was used for substrate thickness, Stokes Radii and TGF-β treatment experiments. One-way ANOVA was used for cellular vs acellular comparisons. Bonferroni’s correction was used for all post tests.

